# Irrigation water depths and soil covers in carrot crop

**DOI:** 10.1101/2020.05.13.094128

**Authors:** Joslanny Higino Vieira, Catariny Cabral Aleman, Elis Marina de Freitas, Laylton de Albuquerque Santos, Gustavo Henrique da Silva, Pedro Henrique Franco Fernandes

## Abstract

The use of soil covers may decrease water consumption and improve the crop sustainability Thus, the objective of this work was to evaluate biophysical parameters and yield of carrot crops (cultivar Brasília) grown under different irrigation water depths and soil cover conditions. The experiment was conducted at the Federal University of Viçosa, MG, Brazil. A randomized block design with four replications was used, in a 5×3 factorial arrangement.The treatments consisted of 5 irrigation water depths, based on the actual soil water capacity (20, 40, 60, 80, and 100% ASWC) and 3 soil covers (white polyethylene, biodegradable semi-kraft paper, and no soil cover - Control). The soil and leaf temperatures, number of leaves, root length, normalized difference vegetation index (NDVI), and fresh root weight were evaluated. The data were subjected to analysis of variance through the F test and the means compared by the Tukey’s test (*p*≤0.05); regression analysis was carried out using the equation with the highest significant fit. The use of semi-kraft paper was a good option for the carrot crop; and the water depths of up to 60% ASWC did not hinder the crop.

## Introduction

Carrot (*Daucus carota* L.) is one of the most important vegetables in the world due to its large planted area and possibility of improving the socioeconomical development of farmers (1) since the return on invested capital is relatively quick. In addition, the root of this plant species has high nutritional value; it contains vitamins, fibers, minerals, and other phytochemicals that are benefic to human health and can prevent cancers (2)(3)(4).

Brazil is the fifth largest carrot producing country (5) where this crop is one of the four more important vegetables. The carrot world production in 2017 was of 42,831,598 Mg (6). Brazil reached yields of approximately 14,077 kg ha in the 2018 crop season, with emphasis on the state of Minas Gerais (7).

Irrigation for carrot crops, as for most vegetables, may increase crop yield and improve product quality, and is an important factor that determines the final production (5). Moreover, the application of an adequate quantity of water, which supplies the water needs of the crop deceases the spread of pests, diseases, and leaching of nutrients (8). Thus, irrigation planning and management should be adopted to obtain the maximum efficiency and preserve the high economical value of carrots in the market (9).

Soil covers can be used to reduce irrigation water depth and decrease the evapotranspiration demand of agricultural crops. Mulching with organic or inorganic materials cover the soil and form a physical barrier that limits soil water evaporation, maintains soil structure, and protects crops against contaminations due to contact with the soil (10).

The soil coverage with polyethylene film is the most used mulching by vegetable producers due to the relatively low price of polymer materials. However, several paper types with potential for use as soil cover can be used, such as semi-kraft, recycled, newspaper paper, and sugarcane straw-based paper (12). Therefore, studies related to this technique are needed to increase the sustainability of vegetable production. The objective of this work was to evaluate biophysical parameters and yield of carrot crops (cultivar Brasília) grown under different irrigation water depths and soil cover conditions.

## Materials and methods

### Experimental site

The experiment was conducted at the Irrigation and Drainage Experimental Area of the Department of Agricultural Engineering of the Federal University of Viçosa (DEA/UFV), in Viçosa, MG, Brazil (20°45’S, 42°52’W, and altitude of 648 m). The experiment was carried out from August to November 2019.

The climate of region is Cwa, tropical highland, according to the Köppen classification, with a rainy season from October to May, and a dry season from June to September. The predominant soil in the experimental area is Typic Hapludult, whose granulometric distribution was 46% clay, 39% sand, and 15% silt. The soil presented density of 1.3 g cm^-3^ in the 0-0.20 and 0.20-0.40 m layers, field capacity of 36.8%, and permanent wilting point of 19.5%, according to the soil water retention curve of the area.

### Plant material and experimental design

A randomized block experimental design with four replications was used, in 5×3 factorial arrangement, totaling 60 experimental plots. The treatments consisted of 5 irrigation water depths, based on the actual soil water capacity (20%, 40%, 60%, 80%, and 100% ASWC) and 3 soil covers (white polyethylene, biodegradable semi-kraft paper, and no soil cover - Control).

Five beds of 12.0 m length, 1.0 m width, and 0.3 m height were raised in the experimental area. Soil fertilization was carried out based on the soil chemical analysis, using the fifth approach for lime and fertilizers uses for the state of Minas Gerais (13)

The carrot cultivar used (Brasília) has high resistance to leaf blight. The plant spacing used was 0.30 m between rows, and 0.10 m between plants; the sowing was done using 3 seeds per pit at 0.10 m depth. A thinning was carried out after the establishment of the crop with cuts at 0.10 m height.

### Irrigation management and Water consumption

The irrigation water depths used were based on the actual soil water capacity, namely 20%, 40%, 60%, 80% and 100%(control).A drip irrigation system was used. The uniformity of the distribution of water by the emitters in field conditions was evaluated. The irrigation management was based on tensiometers installed in the 0-0.10 and 0.10-0.30 m soil layers. The readings of the tensions were converted into soil moisture through the soil water retention curve, obtained by the Richard’s chamber method. The Van Genuchten model (14) was adopted. The irrigation water depths were calculated for each treatment according to Equation 1.

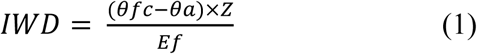

where *IWD* is the irrigation water depth (mm), *θfc* is the volumetric soil moisture at field capacity (m^3^ m^-3^), *θa* is the actual volumetric soil moisture before irrigation (m^3^ m^-3^), *Z* is the irrigated soil layer (mm), and *Ef* is the efficiency of the irrigation system in the test field (0.99).

The water consumption during the carrot crop cycle was calculated based on the Equation 2 of the soil water balance (15).

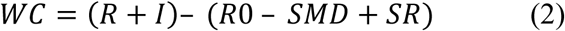

where *WC* is the water consumption (mm), *R* is the rainfall depth (mm), *I* is the irrigation water depth applied (mm), *R0* is capillary increase (mm), *SMD* is the soil moisture depletion (mm), and *SR* is the surface runoff (mm).

In this method, the soil moisture depletion between sampling intervals was calculated by integrating the irrigation water depth applied and rainfall depth between intervals during the crop period. The capillary increase was considered null because the water table was at least 15 m below the ground; soil moisture runoff was also assumed as insignificant because the experimental area is plain.

### Soil temperature

The maximum and minimum temperatures and thermal amplitude of the soil were evaluated daily, after thinning of the carrot plants. Digital sensors (DS18B20) with precision of 0.5 °C from -10 to +85°C were previously installed at the center of the 15 experimental plots to collect temperature data every 30 minutes and store in dataloggers.

### Agronomic variables

Five plants of the central row of each experimental unit were used for the evaluations. The number of leaves was manually counted. Leaf temperature (K) was estimated using an infrared digital thermometer (ST600; Incoterm^®^, Porto Alegre, Brazil) with precision of ± 275.0 K and resolution of 273.1 K. The device was positioned at 15 cm distant from the leaves during the readings.

The length and diameter of the carrots were measured at harvest using a ruler and caliper (mm), respectively. The root fresh weight was evaluated using an analytical balance with precision of 0.01 g.

### Data analysis

The data were subjected to analysis of variance through the F test, using the Experiment Designer package in the R program, and the means were compared by the Tukey’s test (*p*≤0.05); regression analysis was carried out using the equation with the highest significant fit.

## Results and discussion

The water consumption during the carrot crop cycle in the five irrigation water depths and three soil cover types is shown in Fig 1. The carrot crops presented higher water consumption when grown in soil without cover, regardless of the irrigation water depths.

**Fig 1.**
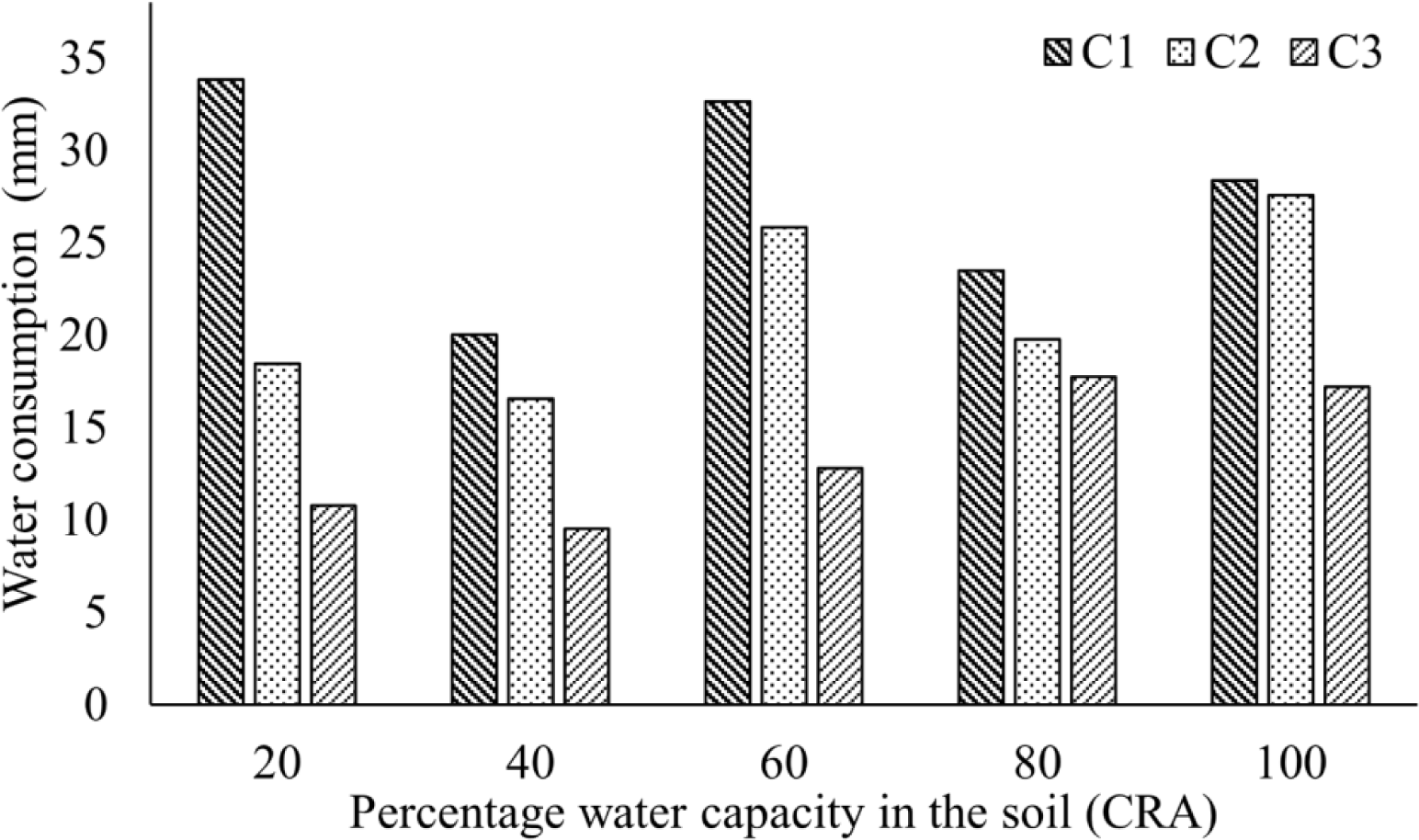
Water consumption by carrot crops (mm) under different percentages of actual soil water capacity (ASWC) and soil covers. Water depths based on the ASWC: L1 = 20%, L2 = 40%, L3 = 60%, L4 = 80%, L5= 100% (control); C1 = no soil cover (Control); C2 = soil with white polyethylene mulching; C3 = soil with semi-kraft paper cover

The highest water deficit used was 20% ASWC (L1), which resulted in a higher water consumption (33 mm) under in the soil with no cover. The water consumption for the white polyethylene and semi-kraft paper soil covers represented savings in water consumption of 45% and 68% when compared to the Control. This was probably due to the soil cover, which decreased soil evaporation rates and increased soil water retention (16).

The treatments polyethylene and semi-kraft paper required lower volume of water over the crop cycle than the Control with no soil cover. This generated decreases in the amount of water applied to the crop. The use of soil cover can generate water savings during the crop cycles of vegetables (17; 12).

The soil coverage with polyethylene resulted in higher crop water consumption than the soil coverage with semi-kraft paper. This may be related to the higher capacity of the polyethylene cover in convert solar radiation, which increases soil heat flux (18), and, consequently, the soil temperature and plant transpiration, which will require more water (19). Soil cover with paper is efficient for water savings, comparable to polyethylene film, and therefore can be used as an adequate alternative to substitute it (20).

The soil maximum and minimum temperatures at 0.05 m depth were lower for the soil coverage with recyclable semi-kraft paper, when compared to the other treatments (Table 1).

**Table 1.**
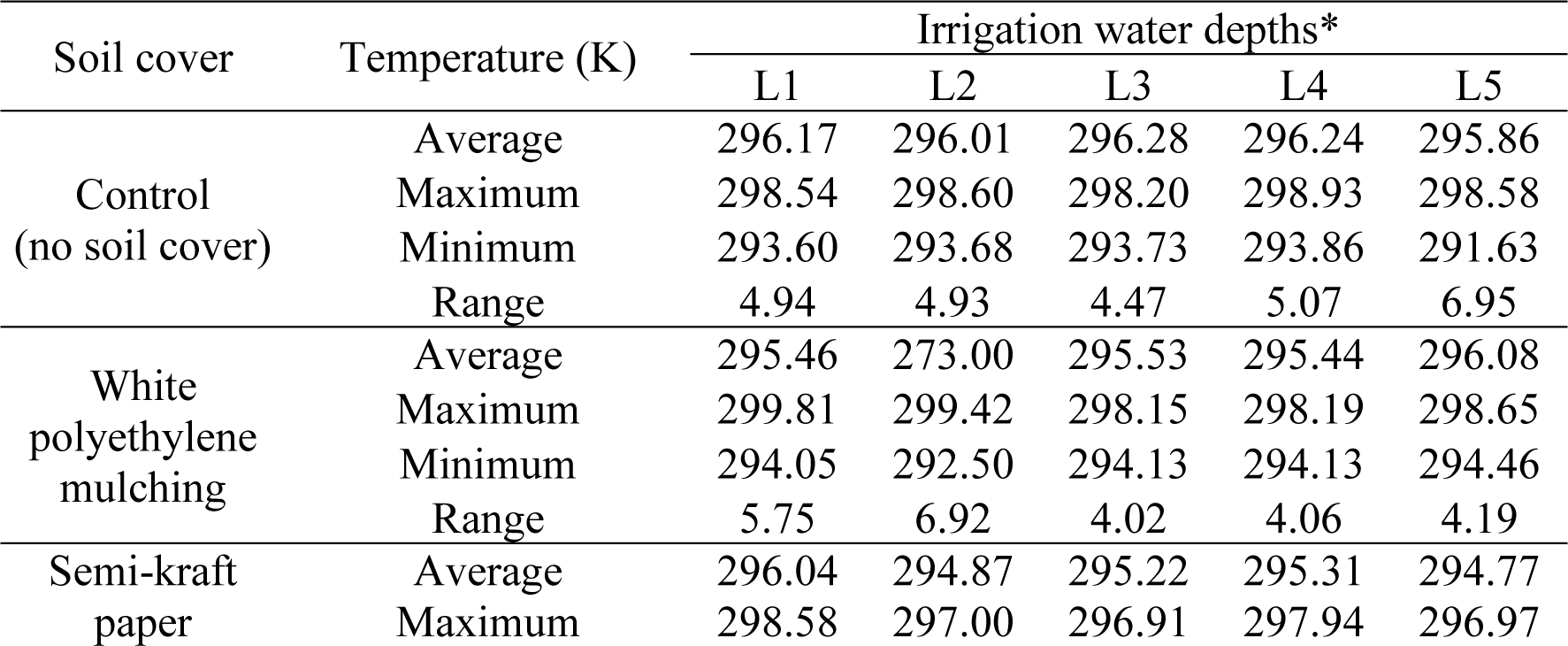
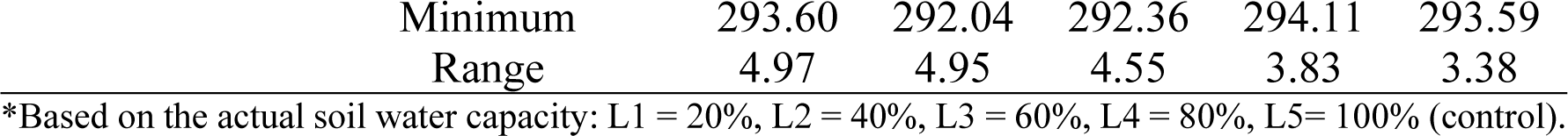
Temperatures at 0.05 m of depth of soils under with carrot crops under different percentages of actual soil water capacity and soil covers.

Table 1 shows that the Control treatment had higher temperatures in all irrigation water depths applied. However, the maximum and minimum temperatures varied according to the water depth used in each treatment.

The highest temperatures were found for the irrigation water depths of 20% and 40% ASWC combined with the soil coverage with white polyethylene, and the lowest temperatures were found for the irrigation water depths of 40% and 60% ASWC combined with the soil coverage with semi-kraft paper. These results were due to the white polyethylene cover, which increases considerably the soil heat flux and, therefore, the soil temperature, while the paper had not the same effect (21).

According to (19), this is because the coverage with white polyethylene considerably increases the heat flow in the soil, therefore, it increases the temperature of the soil, while the paper is not developed in the same way. Similar results were reported for lettuce crops by (22). (23) who found lower soil temperature when using kraft paper as soil cover, when compared to soils under black polyethylene film cover and soils without cover (24) Soils under polyethylene film present mean temperatures of 275,15K and 276,15K higher than those without cover. An experiment with sweet pepper crops showed that soils covered with polyethylene film have temperatures approximately 275,15 K higher than those under conventional hoeing system (25).

Experiments in Japan (26), the United States (27), Vietnam (28) and Mexico (29), also showed higher soil temperatures for soils covered with black polyethylene film. The soil cover can alter the effects of soil temperature, moisture, wind, solar radiation, and evaporation process (30). Covered soils are not affected by the direct incidence of wind and solar radiation, main responsible factors for water evaporation (31).

The use of soil cover reduces soil temperature oscillations and evaporation rates, accelerating the crop development during the initial growth stage, and promote weed control (18), (32) and (33).

The soil without cover had mean temperature of 296 K, with minimum of approximately 293 K. This was due to the soil higher conversion of solar radiation into latent heat due to the lack of physical barrier for soil water evaporation. Thus, the soil temperatures did not decrease as in the other environments evaluated, and were high throughout the crop cycle, with little variation in thermal amplitude. Higher variations in thermal amplitude were found for soils with polyethylene cover when compared to organic material(32).

The agronomic variables root length, root diameter, root fresh weight, and normalized difference vegetation index (NDVI) were affected (*p*<0.05) by the soil covers used (Table 2).

**Table 2.** Agronomic variables of carrot plants (*Daucus carota* L.) grown in soils under different covers

| Cover | Root length<br>(cm) | Fresh weight<br>(t ha<sup>-1</sup>) | NDVI | Diameter<br>(cm) |
| --- | --- | --- | --- | --- |
| Control test | 21,95 a | 21,501 b | 0,68 a | 2.29 c |
| White polyethylene | 19,43 b | 26,265 ab | 0,51 b | 2.78 b |
| Semi kraft paper | 20,83 a | 28,353 a | 0,62 a | 3.20 a |
| Significance level | ** | ** | ** | ** |
| CV (%) | 8,74 | 26,06 | 15,95 | 7,3 |

Means followed by the same letters in the columns are not different by the Tukey's test at \* ( $P\leq 0.05$ ), \*\* ( $P\leq 0.01$ ), ns (not significant)

The root lengths were higher when using soil without cover, but with no significant difference from the soil coverage with semi-kraft paper. The polyethylene cover resulted in lower root lengths, which can be due to decreases of the harming effect weeds by the soil covers (34;35) thus, the carrots grown without soil cover probably needed to compete for water and nutrientes.

The root fresh weight was lower when the crop was grown in soils without cover, with mean of 21.501 Mg ha^-1^; it was better for the crop with kraft paper cover, but with no difference between treatments. However, agronomically, the semi-kraft paper resulted in better means. Many studies show increases in plant growth and gain yield for plants grown in soils under plastic covers (26). A study on water deficits and soil covers for onion crops showed that black polyethylene cover improves, significantly, the water use efficiency (36).

Root diameter is the most important characteristics of carrots for consumers. The soil covers (Table 2) and irrigation water depths (Fig 1) used affected the root diameter. The soil coverage with semi-kraft paper showed the best means, reaching 3,20 cm, corroborating with (37).

The results of the root diameter as a function of irrigation water depths fitted to a quadratic model, with increases up to the highest water depth used (Fig 1). Thus, the highest water depth of 100% ASWC resulted in the highest root diameter; this was due to the balanced environment (soil-plant-water-atmosphere) provided by the water content at field capacity.

**Fig 2.**
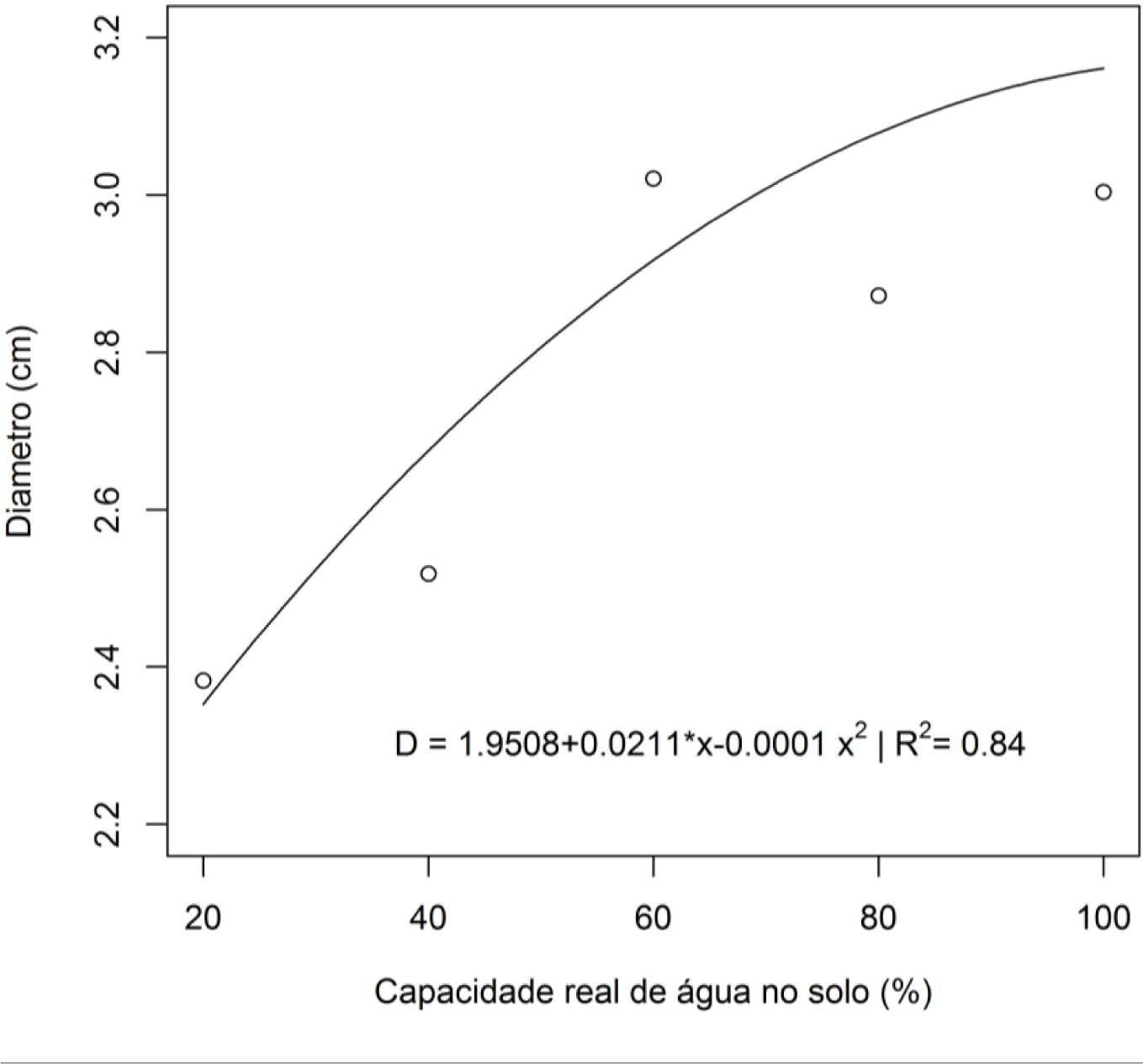
Carrot (*Daucus carota* L.) root diameter under different irrigation water depths. The Normalized Difference Vegetation Index (NDVI NDVI indicates the nutritional state of the plants in relation to nitrogen (38) Higher NDVI was found for the Control treatment with no soil cover, and the semi-kraft paper (Table 2); the higher the NDVI, the higher the crop growth vigor (39). This result indicates a difference in reflectance between the polyethylene and the other cover treatments. In addition, the NDVI was affected by the irrigation water depths (Fig 2), with decreases as the soil water percentages were increased, with a critical point in 76% of ASWC, allowing a maximum point of 0.67.

**Fig 3.**
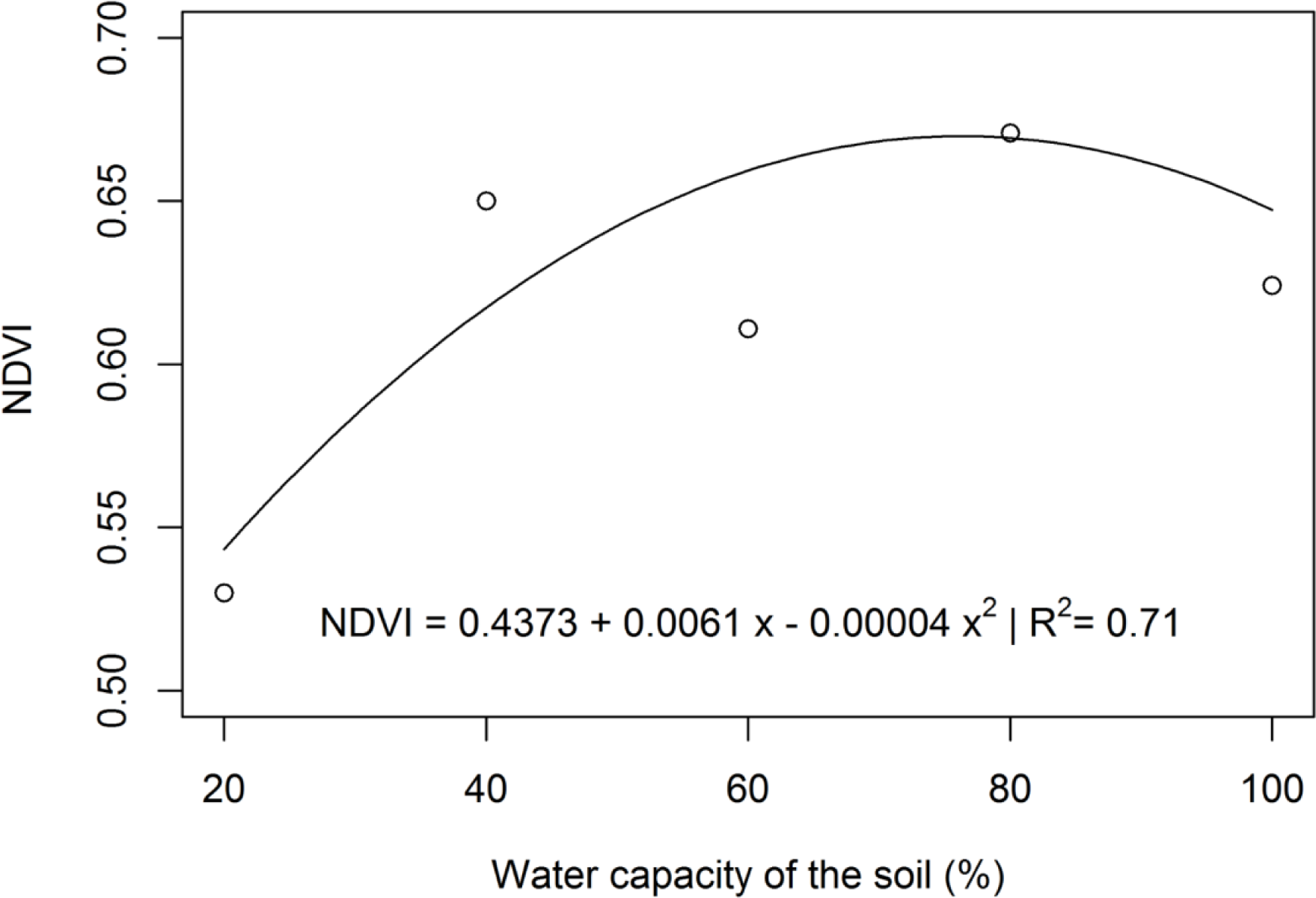
Vegetation Index of Normalized Differences (NDVI) in carrots (*Daucus carota* L.) under different irrigation water depths. The number of leaves and the leaf temperature (Figs 3 and 4) were affected (*p*<0.05) by the irrigation water depths. The carrot market includes a niche for the leaves of these plants. Considering the results of this experiment, the maximum number of leaves can be reached when growing carrot crops under an irrigation water depth of 83% ASWC. These results are different from those of other works (40), which show similar number of leaves per plant for different irrigation treatments.

**Fig 4.**
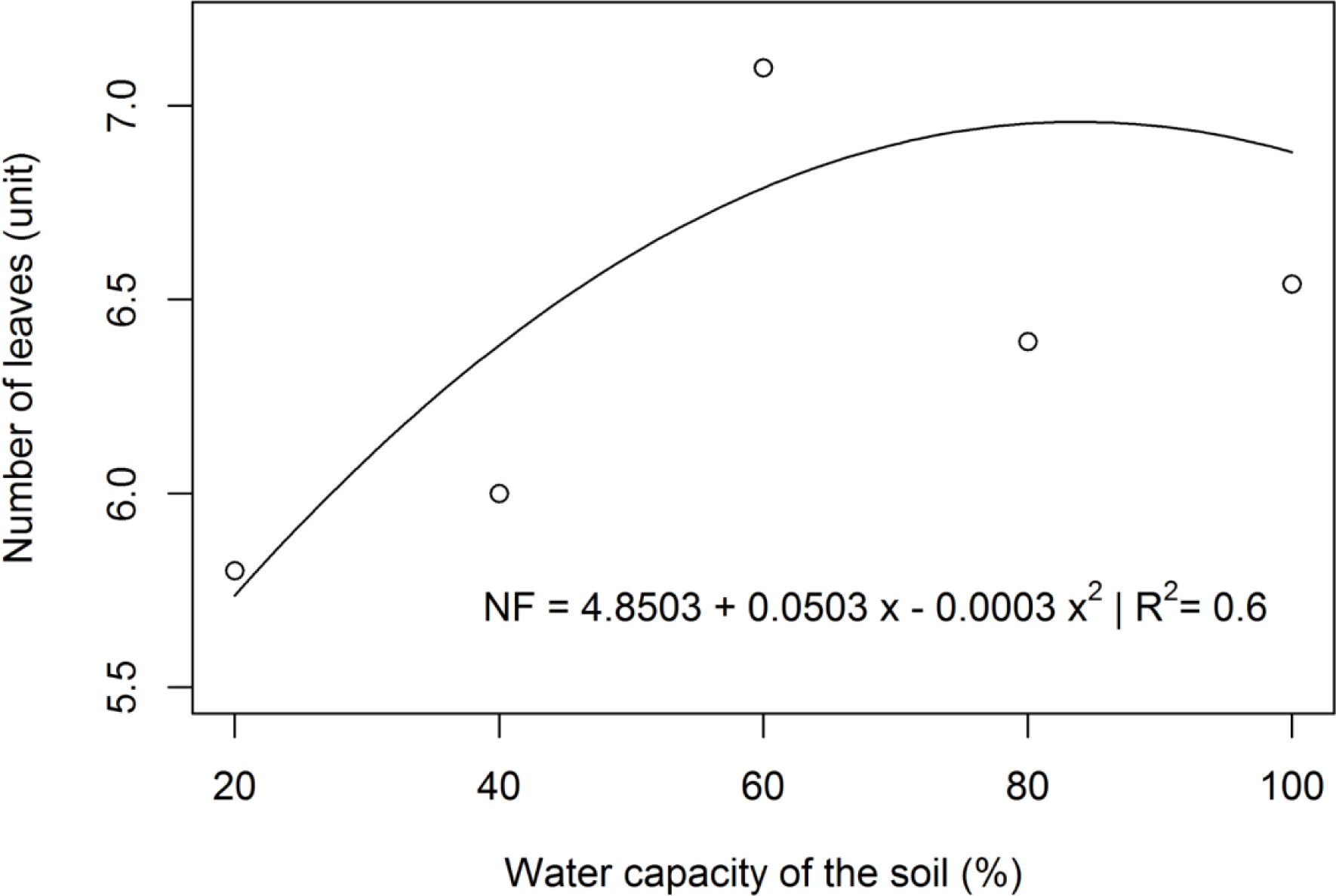
Number of leaves in carrots (*Daucus carota* L.) under different irrigation water depths. The leaf temperature varied little, showing approximately 291.5 to 293 K, and was lower when the plants were grown under lower irrigation water depths (Fig 4). This result may be due to the lower activity of photosynthesis and transpiration processes caused by the limited water to the crop; thus, the maximum carbon assimilation rate can be inhibited by decreases in stomatal conductance and leaf temperature (41).

**Fig 5.**
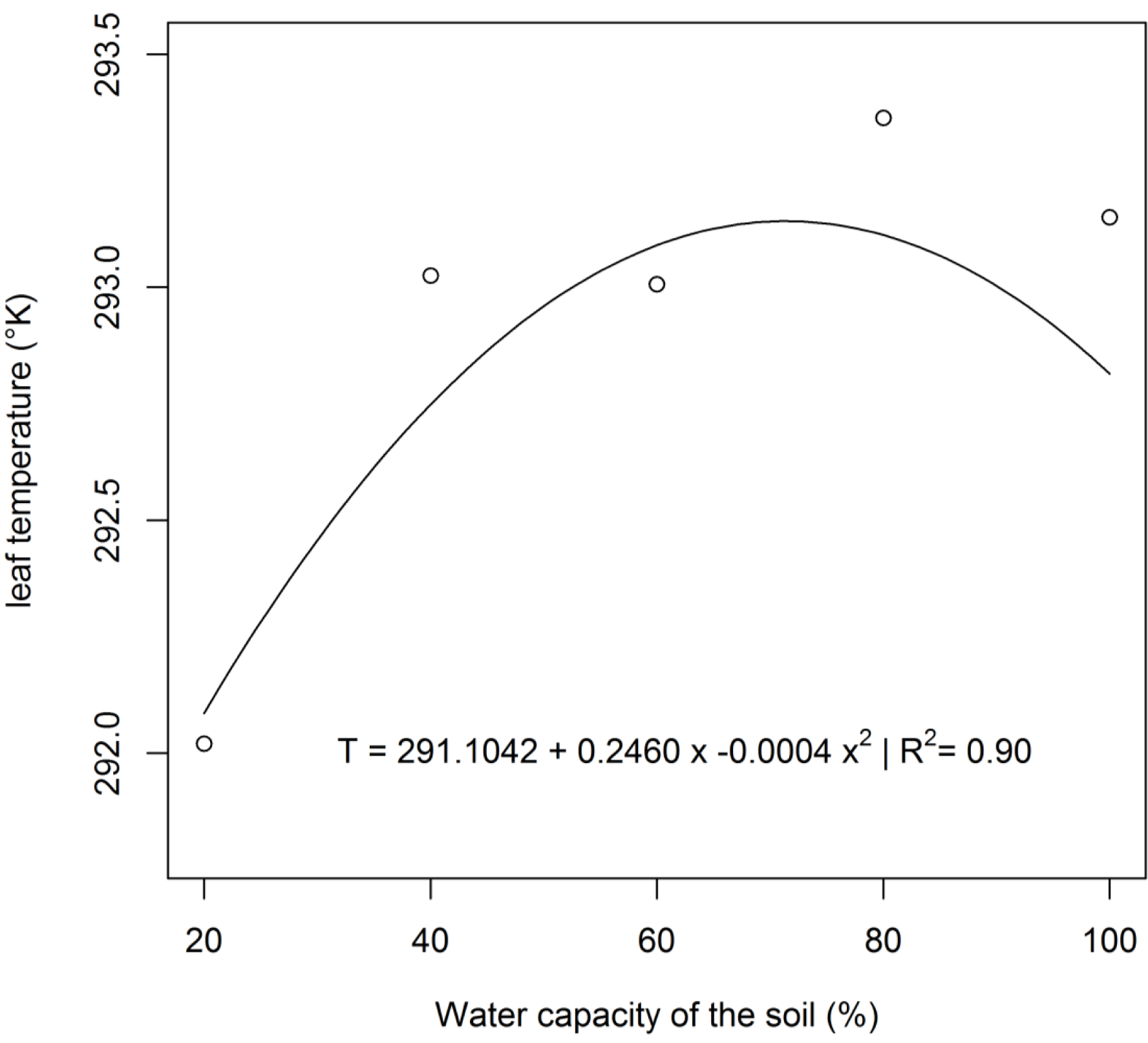
Leaf temperature in carrots (*Daucus carota* L.) under different irrigation water depths. Leaf temperature determines the concentration or water vapor pressure inside the leaf and, therefore, the motor force of transpiration (42). The difference in leaf temperature between plants with and without water stress is due to their water status, stomatal dynamics, and loss of latent heat by transpiration. However, the whole process varies for each plant species, depending on the water stress intensity and duration (43).

## Conclusions

The irrigation water depth of 60% of the actual soil water capacity can be used for carrot production to promote water savings and a good development of the crop.

The use of soil cover with semi-kraft paper is recommended for carrot crops; it provides thermal comfort to plants, even under water stress conditions, and promotes the sustainability of the environment.

